# Encoding manual dexterity through modulation of intrinsic alpha band connectivity

**DOI:** 10.1101/2023.06.27.546709

**Authors:** O. Maddaluno, S. Della Penna, A. Pizzuti, M. Spezialetti, M. Corbetta, F. de Pasquale, V. Betti

**Affiliations:** Department of Psychology, Sapienza University of Rome, Rome, Italy; IRCCS Santa Lucia Foundation, Rome, Italy; Department of Neuroscience, Imaging and Clinical Sciences and ITAB - Institute of Advanced Biomedical Technologies, "G. d’Annunzio" University of Chieti and Pescara, Chieti, Italy; Faculty of Veterinary Medicine, University of Teramo, Teramo, Italy; Department of Neuroscience and Padova Neuroscience Center, University of Padua, Padua, Italy; Department of Neurology, Radiology, and Anatomy and Neurobiology, Washington University, St. Louis, Missouri 63101.

**Author notes:** now, at a different institution (A. Pizzuti). now, at Department of Information Engineering, Computer Science and Mathematics, University of L’Aquila (M. Spezialetti).

## Abstract

Using hands proficiently implies consolidated motor skills, yet malleable to task demands. How the brain realizes this balance between stability and flexibility is unknown. At rest, in absence of overt input or behavior, the communication within the brain may represent a neural *prior* of stored memories. This magnetoencephalography study addresses how the modulation of such stable connectivity, induced by motor tasks, relates to proficient behavior. To this aim, we estimated functional connectivity from 51 participants of the Human Connectome Project during rest and finger tapping in alpha and beta bands. We identified two groups of participants characterized by opposite patterns of connectivity strength and topology. *High and low performers* showed distributed decreases and increases of connectivity, respectively. However, while dexterous individuals also show modulations of the motor network, *low performers* exhibited a stability of such connections. Furthermore, in dexterous individuals, an increased segregation was observed through an increment of network modularity and decrease of nodal centrality. Instead, *low performers* show a dysfunctional increased integration. Our findings reveal that the balance between stability and flexibility is not fixed; rather it constrains proficient behavior.

## Introduction

Healthy individuals differ in their proficiency in using the hand (manual dexterity). One of the critical aspects underlying skilled manual behavior is flexibility, *i.e.,* the capacity to expand the motor repertoire ^1^. This adaptability must reconcile with the requirement of stability to prevent the degeneration of the acquired skills. This inherent tension between flexibility and stability is also crucial at the level of neural circuits. The brain is modified through experience and learning to form long-lasting memories, yet it retains past motor memories ^2^. However, this trade-off is not fixed because of individual differences. One of the primary sources of such a variability is prior learning, *e.g.,* how we learn to execute a task constraints successive motor strategies ^3^. Despite evidence suggesting that prolonged manual practices and motor learning produce brain structural and functional changes ^4–8^, little is known on how manual dexterity and interindividual motor variability are encoded in the flexibility and stability of the communication in the brain.

Interestingly, even at rest, *i.e.,* in the absence of overt input or behavior, the architecture of the interaction between brain regions closely resembles those observed during a motor task ^9, 10^. One leading hypothesis is that resting state networks (RSNs) are sculpted over the lifespan of the individual through previous experiences and learning ^6, 11, 12 13^. These neuroplastic effects can persist up to a week after learning ^12^ and seem to be related to off-line memory processing or motor skills retention ^14, 15^. This is of fundamental importance since procedural memories must be stable and long-term. Therefore, the intrinsic activity is a putative mechanism for preserving motor (and cognitive) representations. In fact, being its topography robust across participants and recording sessions ^16^ and resilient across behavioral states and levels of consciousness ^17–22^, it satisfies the stability requirement. On the other hand, performing a task modifies the strength of the intrinsic functional connectivity ^18–21^. This malleability meets the requirement of flexibility.

Flexibility is a fundamental mechanism allowing adaptive behavior and supporting learning ^23^. It can be realized through modulations of segregation (*i.e.*, independent processing in specialized networks) and integration (*i.e.*, communication among networks) permitting a flexible reconfiguration of the system ^23–25^. Crucially, a dynamic balance of segregation and integration has been proven to support behavioral performance ^26, 27^. Learning processes (*e.g.*, motor learning; ^24^) further modulate the underlying topology, *e.g.,* modularity (decomposability into distinct partitions of the network) and hub centrality (the role of highly interconnected regions). Here, we aimed to explore whether manual dexterity corresponds to different patterns of stability or flexibility of the underlying topology of the brain networks, observed during a motor task or at rest. Built on this evidence, this Magnetoencephalography (MEG) study explored changes of the intrinsic large-scale connectivity (*e.g.*, strength, frequency content) and architecture (*e.g.* integration/segregation) induced by a manual task (or foot movements as a control condition, see *Materials and Methods*), to test if the flexibility and stability of intrinsic architecture encodes the manual dexterity. To this aim, MEG connectivity was measured as synchronous modulations of band-limited power (BLP) ^28–31^ at rest and during the motor tasks in alpha (α) and beta (β) bands. We already showed that connectivity modulations are found in regions involved in the specific task and correlate with performance^25^. Our hypothesis is that the hand movement triggers small modulations mainly involving the motor network, occurring over a backbone topography remaining stable across rest and task states and that preserves stored skills. However, we expect that the specific patterns of functional reorganizations, *e.g.,* modularity and nodal centrality, vary with the proficiency in using the hand. Our results reveal a direct link between interindividual differences in manual dexterity and specific patterns of stability/flexibility of the large-scale network connections, specifically in the α band.

## Results

We analyzed Magnetoencephalography (MEG) data of 51 healthy participants from the Human Connectome Project (HCP) S1200 Release. Participants fixated a small cross (resting-state) or performed a motor task (*i.e.*, finger tapping and toe squeezing) (Fig. 1A) ^32^. After removing task-evoked activity, we computed the leakage-corrected band-limited power (BLP) from the source-space signals in the frequency bands alpha (α, 8-15 Hz), low beta (β, 15-26 Hz) and high β (26-35 Hz) (see *Materials and Methods*). We used a functional parcellation with 164 nodes and 10 RSNs (Fig. 1B) to estimate functional connectivity (FC) from BLP in each subject, condition, and band (Fig. 1C). Then, we adopt a linear model to study the modulation of intrinsic functional connectivity induced by the motor task (Fig. 1D). Based on these results, for the hand condition, participants were clustered. The topological aspects of segregation/integration of the obtained classes were investigated through graph measures such as the modularity and centrality (Fig. 1E). We used the Nine-Hole peg test scores (provided by the HCP) for assessing finger dexterity ^33^.

**Figure 1.**
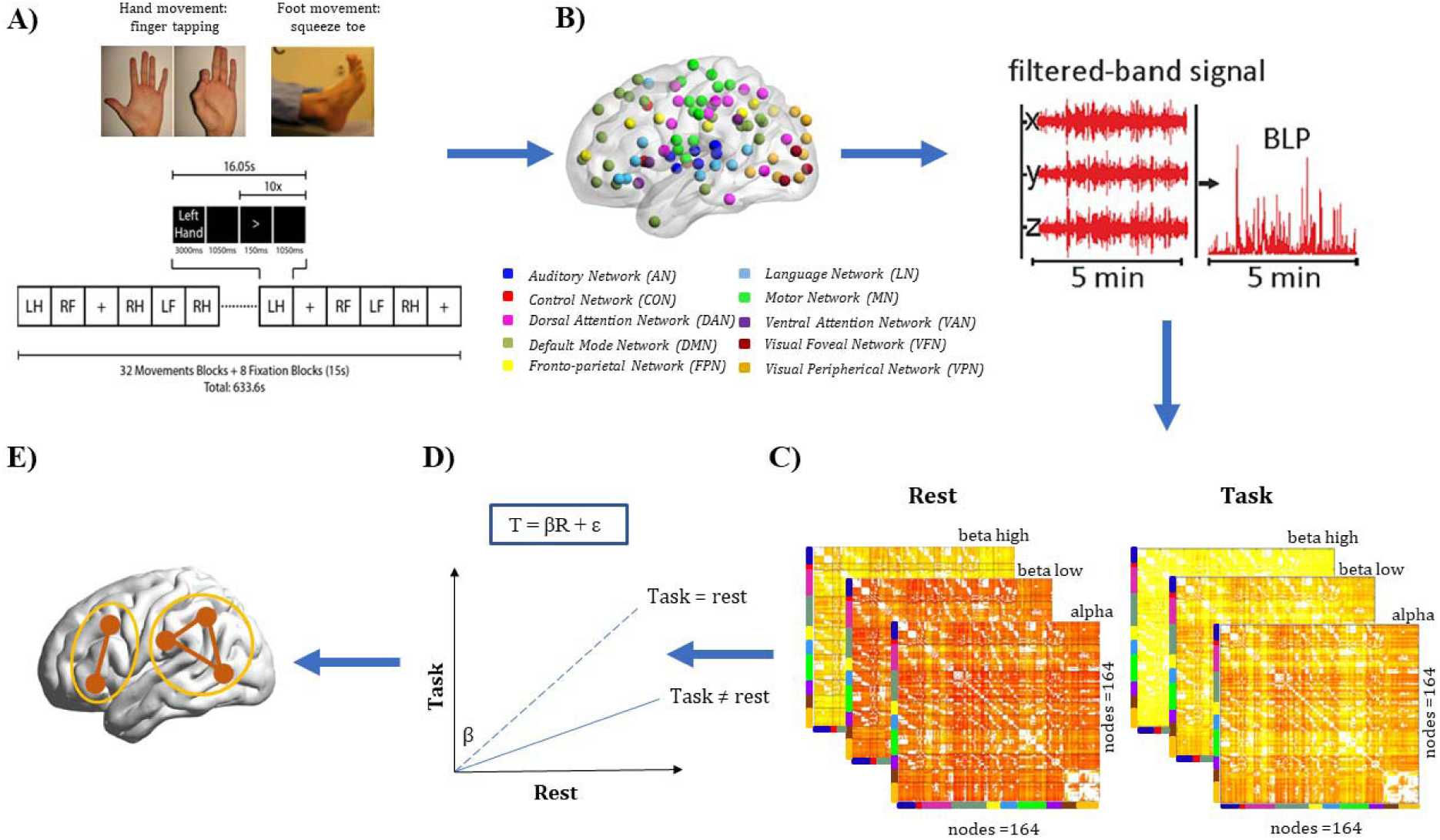
Experimental paradigm and analysis pipeline. **A)** Participants performed a finger tapping (with their right or left hand) or a toe squeezing (with their right or left foot). The motor task consisted of 32 movement blocks (16 hand and 16 foot). Each block consists of 10 trials. 3 resting state runs lasting 6 minutes each precedes the motor task. **B)** A set of 164 node brain parcellation, comprising 10 networks, is used to estimate the source space Band-limited power (BLP) in the alpha, beta low and beta high band. **C)** The static functional connectivity is calculated as the leakage-corrected correlation between each pair of nodes, separately for the resting state and the task data. **D)** Linear model of the relationship between rest and task. A k-means algorithm on the beta values of the model identifies two groups (*i.e.*, *high* and *low performers*). **E)** Measures of segregation/integration measures are computed.

### Hand movements decrease the strength of functional connectivity

In analogy with previous studies ^18, 20–22^, we ascertained that the execution of motor tasks preserves the overall topography of MEG connectivity across all RSNs. The observed stability (Mantel test, p<0.05, r>0.86) of resting-state FC patterns during the motor task suggests a similar functional architecture underlying these conditions (Figure S1). Next, we investigated the influence of hand movements on the intrinsic FC as compared to a control condition (foot movements). We ran repeated measures ANOVA with band (α, low β, high β), condition (hand-rest vs foot-rest), and network (all RSNs) as within-subject factors on the connectivity modulation (task-rest) averaged across connections of each RSN. The movement of the hand (Fig. 2A) and foot (Fig. S2A) induced an overall decrement of connectivity across all RSNs, condition and bands.

**Figure 2.**
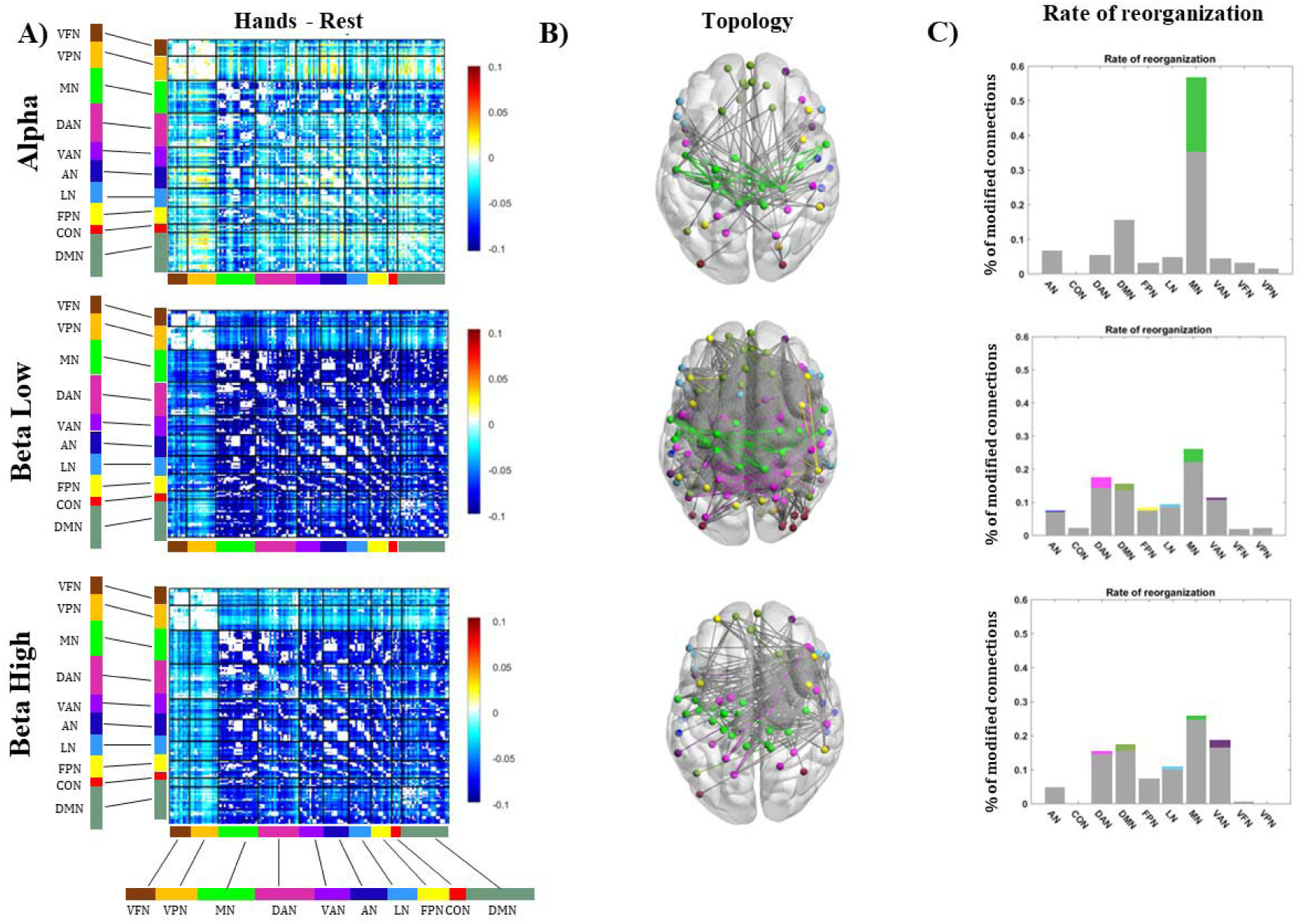
Changes of functional connectivity and topology induced by the finger tapping. **A)** Group level difference connectivity matrices task-rest for alpha, beta low and beta high bands. **B)** changes of network topology. Hands movements modulate fewer links in the alpha band, especially within the motor network (green edges). Conversely, in the beta band there is a wide-spread topological reorganization across all networks. **C)** Percentage of modulated links. Within-network connections are color-coded. In gray between-network connections are shown.

Specifically, we found a significant main effect of band (*F*_2,100_ = 14.90, *p* <0.0001, pη^2^ = 0.23) as accounted by smaller decrements in α as compared to all other bands (mean FC_alpha_ =-1.54, FC_betaLow_ = -3.81, FC_betaHigh_ = -2.59, post hoc Bonferroni corrected, *p*<0.05). We observed also a significant main effect of condition (*F*_1,50_ = 7.01, *p*<0.05, pη^2^ = 0.13) with higher decrements during foot as compared to hand movement (mean FC*_hands_* = -2.31, mean FC*_feet_* = -2.98; *p*_Bonff_ <0.05) (Fig. 2A and S2). Furthermore, we found a significant effect of network (*F*_9,450_ = 13.81, *p*<0.0001, pη^2^ = 0.22) as well as interaction band x network (*F*_18,900_ = 9.19, *p*<0.001, pη^2^ = 0.16) accounted by a smaller decrement in the motor network in the alpha band (FC_alphaMN_ = -1.3) as compared to the beta low (FC_betalowMN_ = -3.81) and beta high bands (FC_betaHighMN_ = -2.72) (*p*_Bonff_<0.001). Finally, all the other interactions were significant (all p-values <0.05) except for band x condition. See SI for details.

To summarize, in agreement with previous MEG reports on visual stimuli ^18, 19^, all RSNs decreased their connectivity during the motor performance, despite maintenance of the overall topography. Functional connectivity decrements spread along the entire cortical mantle. Interestingly, we found a different modulation in the α band compared to the other bands. Crucially, the modulation of the Motor Network in α is significantly different compared to the β bands. Finally, we also found different FC modulations produced by hand vs foot movements.

### Moving the hand reorganizes the network topology

Next, we investigated changes of functional topology induced by the two motor tasks by means of Network-based Statistics (NBS ^34^). Figure 2B shows graph components that significantly decreased for each frequency band for the hand movement. In the α band, the task produced a high proportion of decreased connections, especially in the motor network (Fig. 2B, upper panel, green lines). Differently, in the low β band while a larger number of connections were also reduced, we did not observe a predominant proportion of connections involving the motor network (Fig. 2B middle panel). The same applies to the high β band, where we observed an intermediate number of decreased connections (Fig. 2B lower panel). To quantify this observation, for each RSN, we computed the percentage of modified links, normalized by their total number (Fig. 2C). The bar plots illustrate the between- and within-network connections for the three bands. In the α band, the motor network shows the highest percentage of altered within-network connections, namely 56% (upper panel). Conversely, in the β bands (both high and low beta, middle and lower panels) (Fig. 2C), we found a modulation of between- and within-network topology involving all RSNs, with larger components in the β low (percentage of affected links = 94,8%) and in the β high (percentage of affected links = 92,5%) than the α band. As a control, we performed the same analyses for toe squeezing: in general, foot movement re-organized a large number of connections in all bands. However, we did not observe a specific pattern (Fig. S2B-C).

In summary, hand movements induce a topological reorganization. While this is extensive in both β bands, the motor network predominantly modulates in the α band. These results may suggest different functional roles of the α and β bands in the manual dexterity.

### The modulation of alpha band connectivity encodes manual dexterity

Now, we addressed whether task-based modulations of intrinsic connectivity are a distinctive feature of the hand movement and thus can be linked to manual skills such as dexterity. First, to evaluate the similarity between FC at rest (rFC) and during the task (tFC), we adopted a linear model: *tFC =* β *x rFC + e* (Fig. 1D) and we studied the variations of β values *(i.e.,* the slope). Specifically, for every RSN, we considered its set of connections (both within and across the other RSNs) and from them we estimated the β values. This provides, for each subject and frequency band, a matrix of β, with β*ij* representing the stability between rFC and tFC between network *i* and *j*. Specifically, the larger is the distance of β*ij* from one, the higher is the instability between rFC and tFC). In this way, for every participant, we obtained a specific profile of functional reorganization between task and rest. Then, we clustered subjects based on their β profiles. The optimal number of classes for the clustering algorithm was estimated by applying a data-driven approach (see SI).

For the α band, the optimal number of clusters was two (see SI, Fig. S3). The centroids of the obtained clusters, *i.e.,* the average of the β matrices within each class, are reported in Figure 3A and S4. Specifically, in the first group (Fig. 3A left panel), β values were smaller than 1 (mean value=0.80), suggesting a low stability of the considered connections between task and rest. By contrast, in the other cluster (Fig. 3A right panel), β values were closer to 1 (mean value = 1.08), suggesting a lower flexibility of the overall network. This different trend is evident in the display of the linear fitting over the links and subjects (Figure 3B). Then, we tested whether the two groups reflected different behavioral performance, as investigated through the manual dexterity task (*i.e.*, the Nine-hole peg test). Interestingly, we obtained a significant difference in terms of dexterity (*t*_49_ =-2.94, *p*=0.005). One group (*i.e.*, *high performers*) was characterized by faster reaction times (mean RT = 95.96 ms), while the second one (*i.e.*, *low performers*) showed slower reaction times (mean RT = 103.63 ms) (see Fig. 3C). Notably, this result is specific for the α band. We did not obtain any significant clustering either in low or high β bands (see Table S1).

**Figure 3.**
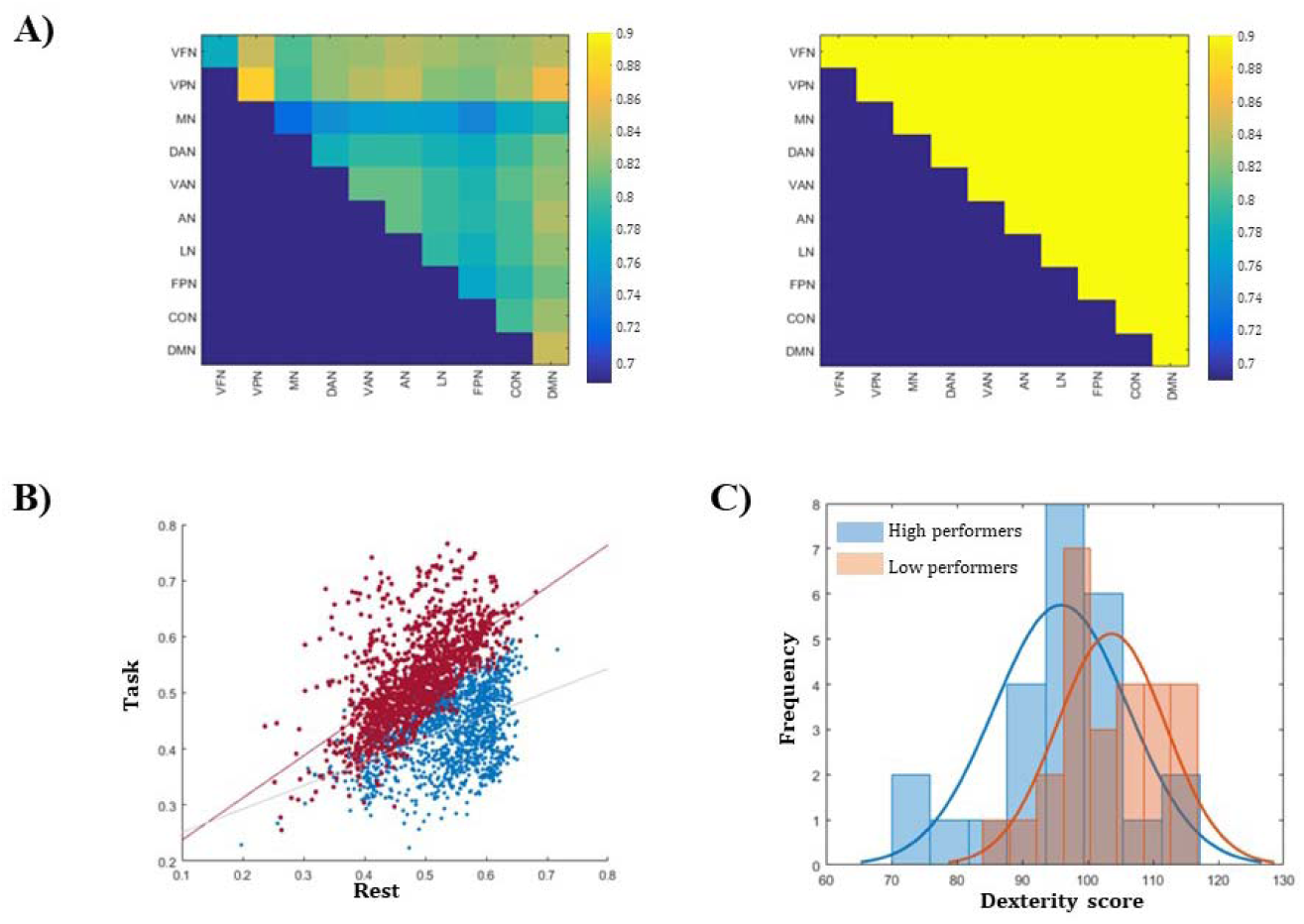
K-means clustering identifies low and high performers. **A)** Centroids of the two clusters, consisting of beta values, identified by K-means. **B)** Scatterplot depicting the linear relationship between rest and task connectivity in the two groups (blue: high performers, red: low performers). A distinct linear trend is evident. **C)** The 2 groups differed in terms of manual dexterity. The performance was measured as mean RTs (see Materials and Methods). The RT histogram is reported for *high* land low *performers*.

Based on these findings, we then tested the stability of the β values and the related clustering when pruning, at increasing levels, the original set of connections, to assess which was the range of connections mainly driving the classification. To do so, we thresholded the connectivity matrices preserving only connections above a N percentile value (see Table S1), from N = 0 to N = 0.99. We then computed again β values for each network subset and we repeated k-means clustering. Our results demonstrate that the algorithm separates two groups differing in dexterity, only for connections not exceeding the 86th percentile. Stronger connections do not contribute to β patterns different across the two groups. Overall, this result suggests an encoding of manual dexterity in the task vs rest modulations of the α-band BLP connectivity.

### High and low performers differ in terms of connectivity

Once we separated participants into two groups (*high* and *low performers*), we analyzed whether they also exhibited different modulations of network topology. First, separately for the two groups, we compared the pairwise rFC and tFC through a paired t-test. As shown in Figure 4A, *high* and *low performers* show two distinct patterns of FC modulation: *high performers* show an overall significant decrease of connectivity across all RSNs. Conversely, l*ow performers* display a general significant increase of FC strength. This does not apply to the Motor Network (followed by the Dorsal Attention Network-DAN and the Visual Foveal Network-VFN), which instead appears relatively unchanged with respect to rest. Second, we tested which graph components were modulated by finger-tapping separately for the two groups. As shown in Figure 4B, participants with higher dexterity show significant decreased components, representing a selective reorganization of between-networks connections. On the contrary, *low performers* show significant increased components, representing changes in both between- and within-network interactions.

**Fig 4.**
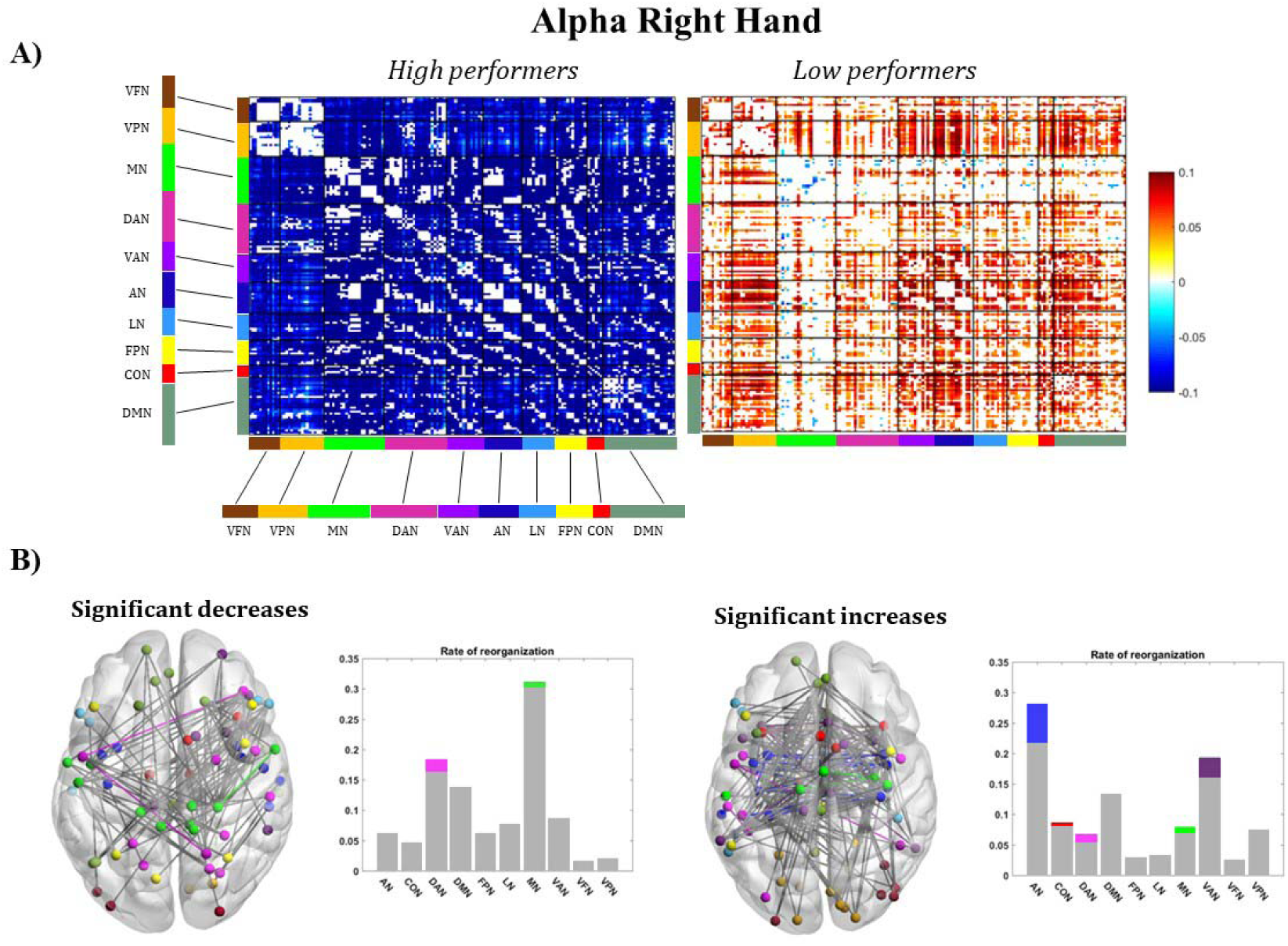
FC and topology changes relate to manual dexterity. **A)** Difference connectivity matrices (task-rest) thresholded after the t-test for *High (left panel) and Low performers (right panel).* In the matrices are depicted only connections with a significant p-value. *High performers* exhibit an overall decrease of FC in all networks, while *low performers* show a slight increase, with a stability in the Motor Network. **B)** Modulation of topology in *high* and *low performers* and percentage of modulated links. Within-network connections are colored-coded. In gray between-network connections are shown.

Based on these results, we next analyzed patterns of connectivity modulations involving the strongest links in each condition. Specifically, in each condition and for each subject, we identified the strongest connections as the ones above the 86^th^ percentile. We then estimated the tFC-rFC difference, and we evaluated the consistency of connections decreasing or increasing across subjects, *i.e.,* the percentage of occurrence of each connection in every group. Notably, *high* and *low performers* exhibited different modulations of the strongest links. The majority (70% of subjects) of *high performers* showed a decrease of connections involving the DMN (Figure 5A, left panel). Notably, this decrease involves exclusively intra and inter DMN connections with no other interactions. This suggests that, among the strongest connections in both conditions, the ones involving the DMN are consistently reduced. As far as eventual increase of communication, we did not observe any connection in this group. At the same consistency level (70%), *low performers* display an increase of interactions involving the strongest links (Fig. 5A, right panel). This consists of a set of task-unrelated networks, involving the DMN but also VPN, AN, VFN and CON. By mapping these connections, we found a precise spatial topography of these changes, following two opposite axes: an antero-posterior stream related to a decrease in communication in *high performers* and an interhemispheric stream linked to an increase of communication in *low performers* (Fig. 5B).

**Figure 5.**
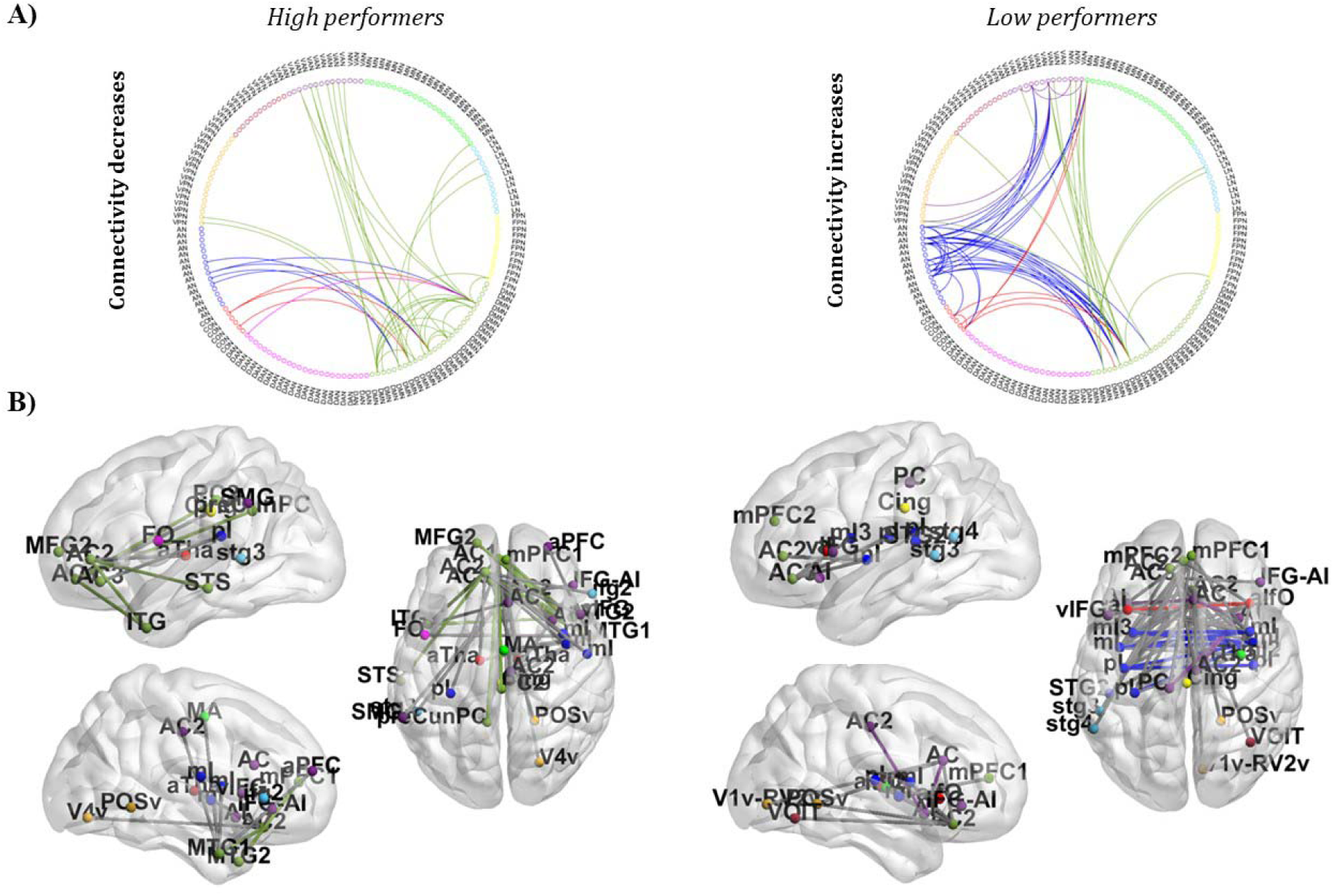
Spatial topography of reorganization in the two groups. **A)** In *high performers* the overall communication involving DMN strongest links decreases (right panel). On the contrary, in low performers, the architecture of strongest links shows an overall increase (left panel). **B)** In *high performers,* the modulation of strongest connections induced by finger tapping involves the DMN along an antero-posterior axis (left panel). In *low performers,* instead, an inter-hemispheric axis, involving task-independent networks (right panel), is involved.

### Task-induced modulations of segregation/integration reflect subjects’ dexterity

Our previous evidence suggests that there are specific patterns of FC reorganization depending on different participants’ motor skills (*i.e.*, different manual dexterity). Previous studies showed that motor learning induces increased segregation of the sensorimotor system and reduction of hub centrality ^24^. Analogously, we expected that a skilled behavior induces changes in network integration/segregation. In the α band, we estimated the graph modularity by quantifying the amount of segregation. To this aim, we first applied a percolation analysis ^35^ to obtain individual binary graphs. Then, we applied the Louvain modularity ^36^ and ran a mixed model ANOVA on modularity, with Group (*high* and *low performers*) as the between-subject factor and experimental condition (rest, motor task) as the within-subjects factor. We found a significant interaction Group x Condition (*F*_1,49_ = 18, *p*<0.001, pη^2^ =0.27): in *high performers*, the network modularity significantly increased from fixation to the finger-tapping task (*p*<0.005); conversely, in l*ow performers*, we showed the opposite trend (*p*=0.007). Moreover, the modularity during tasks in *high performers* was significantly larger than *low performers* (*p*<0.0005) (Figure 6A). Changes in network segregation occurred with modulations of centrality, as measured through the participation index (PI, ^37^). Figure 6B shows, for each RSN, changes of the PI for the two groups, when going from rest to the motor task. Interestingly, *high performers* show a general decrease of the PI, reaching significance in the Control Network (CON; *p*<0.009) and MN (*p*<0.0003). These results were revealed by the post-hoc analysis on the significant interaction (*p*<0.004) obtained from a mixed model ANOVA with Group (*high* and *low performers*) as a categorical factor, experimental condition (rest, motor task), and RSNs (all networks) as factors. Conversely, *low performers* showed a significant increase of the PI in DAN (*p*<0.0008) and MN (*p*<0.01). When comparing the PI modulations, with repeated measures ANOVA with Group (high and low performers) as categorical factor and RSN as factor (significant interaction p< 0,004), we obtained a significant difference across groups for the MN (p=0.015) and a tendency to significance in the DAN (p=0.058), as shown in Fig.6B. At the regional level, these differences were tested node-by-node through a t-test to look for nodes driving PI modulations. At regional level, *high performers* show a significant decrease of centrality in nodes of the MN (rdPoCe, rmdSPL, ldmSPL, lS2), LAN (lstg4), VAN (IFG-AI), FPN (PrCu) and in the DAN (rmIPS, rpIPS-SPLd). Conversely, in *low performers* the motor task induced a significant PI increase in the MN (rCS, ldPoCe, lvPoCe), DAN (rFEF, lvPoCe-SMG, lPrCe) and a decrease in VFN (rLO) (Fig. 6C).

**Fig 6.**
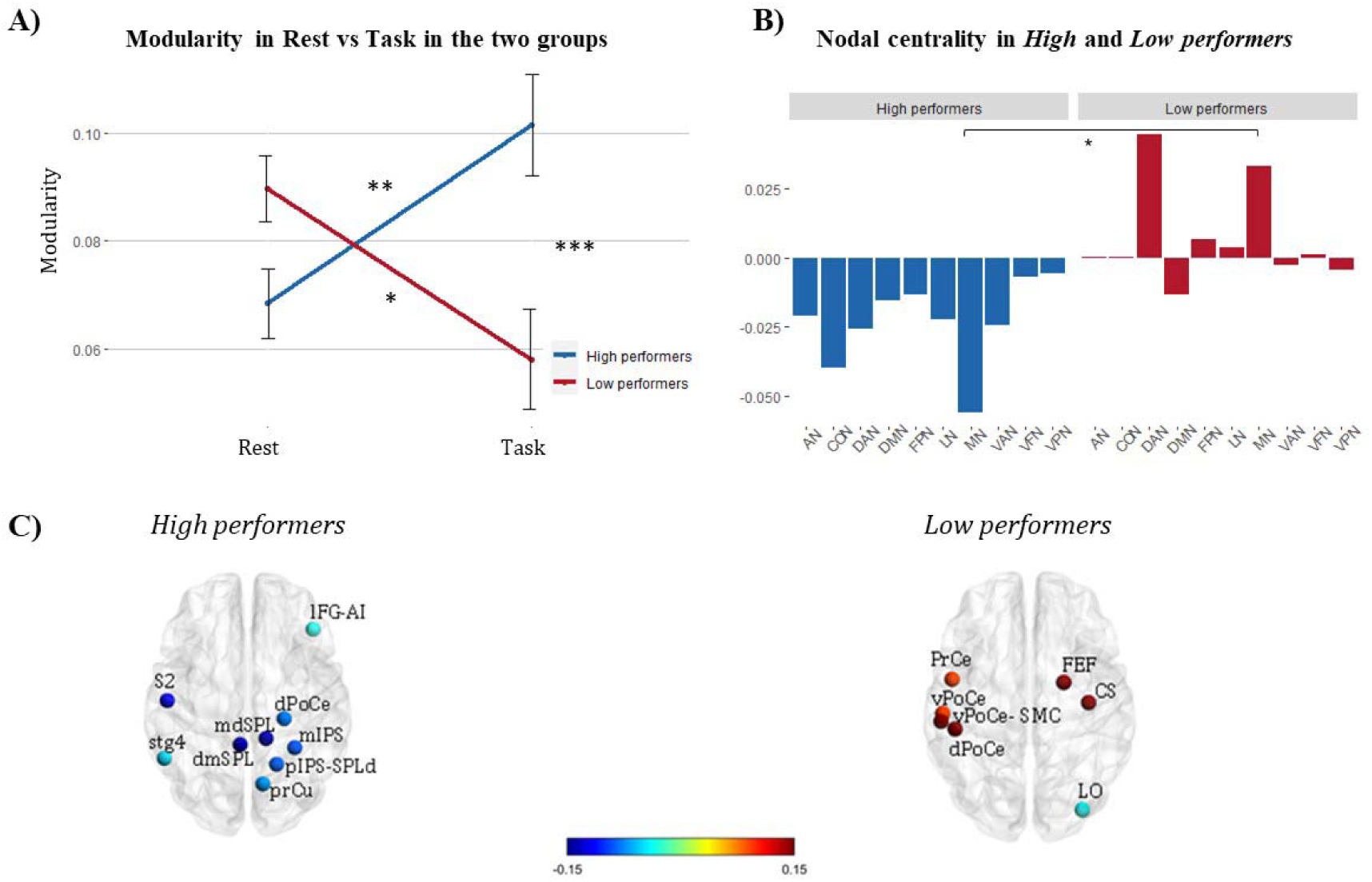
Functional integration and segregation in high/low performers. **A)** *High performers* exhibit higher modularity during finger tapping than rest; conversely, modularity decreases in *low performers* when switching from rest to motor task. It can be noted that the two groups differ in modularity, thus suggesting that segregation relates to the performance. **B)** The participation index shows a decrease of nodal centrality (averaged over nodes in each RSN) in *high performers*, especially in the Motor and Control Network. Conversely, in *low performers,* the participation index increases. This modulation is significant in MN. **C)** Nodes with significant differences in Participation Index between rest and task in *high* (left) and *low performers* (right).

In summary, when executing the finger-tapping task, dexterous individuals show an increase in the segregation of the intrinsic network topology. In parallel, they show a reduction of centrality, mainly in the MN and the CON. Opposite patterns are found in low-performers subjects, with an increase of across-network communication mainly involving the DAN and MN.

## Discussion

Individuals differ in manual dexterity. This study tested whether and how this feature is encoded in the task-induced modulation of spontaneous brain connectivity. First, we confirmed that the topography backbone observed at rest resembles the one during a motor task. However, in the α and β bands, this stability of the network topography combines with a set of consistent changes of connectivity strength and topology. In the α band, a specific reorganization of connections allows to distinguish *high* from *low performers*. This reorganization occurs in opposite directions for the two groups: in *high performers,* it involves mainly a decrease in motor network connections, in *low performers* these connections are more widespread, *i.e.,* not only related to motor network, and either slightly increase or remain more stable. Further, in *high performers*, we observed an “internal focusing” effect in the network topology. This was characterized by an increment of network modularity, thus suggesting an increased segregation paralleled by a decrease of nodal centrality. An opposite trend characterized *low performers*, suggesting a dysfunctional increased integration.

### Alpha band as a marker of manual dexterity

Proficiency in using hands reveals through specific modulations of functional architecture in the α band. The functional significance of this rhythm is multifaceted and under debate. Traditionally, its amplitude has been associated with the inhibition mechanisms of task-irrelevant regions in the brain ^38, 39^. α is an “idling” rhythm ^40^, *i.e.,* denotes a state of inactivity of the brain circuits ready to be modulated during a task. Since more specific task-related patterns of functional connectivity must emerge, the α connectivity needs to be suppressed ^18, 19^. Such an effect (Figure 2; Figure S1) is also in line with decrements of correlated cortical noise occurring during tasks or stimulus presentation, observed in the monkey and cat visual cortex ^41, 42^. At the same time, α oscillations have been linked with high-order cognitive functions, such as memory and attention ^43, 44^. They correlate with faster reaction times, better memory performance and information processing ^45–47^. A link between α rhythm and behavior has been recently highlighted during resting-state, particularly in expert populations. In a sample of pianists, the spontaneous phase coupling in the α band correlates with the motor performance of finger-tapping ^48^. Analogously, in expert dancers, resting-state connectivity successfully decodes the level of motor expertise ^49, 50^. Such an effect has a specific electrophysiological signature, *i.e.*, the mu rhythm ^49^. Overall, these observations suggest a link between the long-term experience, behaviorally relevant increments of spontaneous coupling and specific brain rhythms. This is in line with previous works demonstrating that motor ^6, 11, 12, 51, 52^ and perceptual learning ^53^ shape intrinsic connectivity. Interestingly, resting-state cortical connectivity also predicts motor skill acquisition ^54 55^ and interindividual variability in motor performance ^56^. Here we provide several advancements to these findings.

Although these reorganizations have been mainly observed under laboratory conditions, theoretical models predict that spontaneous connectivity reflects the training of cortical networks occurring during daily life ^57–59^. In this work, we considered a finger tapping task. Now, albeit finger-tapping is a controlled movement, it closely resembles the precision grip between the thumb and index finger ^60, 61^, a most-used hand posture in daily life movements with high consistency across subjects, both at spatial and temporal level ^62^. Thus, one might argue that such a movement, repeated during everyday activities to grasp small objects, is stored into specific patterns of intrinsic connectivity to support motor skills retention. Moreover, our findings show that the proficiency underlying such a movement seems to rely not on the spontaneous connectivity but on its modulation induced by the task. Specifically, the dexterity seems to rely on the direction of this modulation within the involved neural circuits. Specifically, we observed that finger-tapping produces a modulation of the intrinsic communication (both connectivity strength and topology) especially in the motor network, likely supporting the small adjustments required to control the movement. In *high performers*, we found regionally specific decrements in the alpha band. Instead, in low proficient individuals, the motor network does not reorganize and the overall connectome shows increments of the connectivity strength. As a control, during foot squeezing, a less frequent movement not recurrent on a daily basis, we obtained a global reduction of the intrinsic α connectivity. In this case, this reduction is widespread, *e.g.,* it does not involve specific patterns, as in the previous case. Interestingly, this is in line with what we found in the visual system for the observation of synthetic movies (*e.g.,* scambled). We reported stronger changes of the intrinsic connections, as compared to natural movies ^19^. Overall, this first set of results paves the way for a novel mechanistic role of the α-rhythm and its relationship with behavior. The suppression of α-connectivity, well established during a task, seems to be sculpted by the experience. Within this suppression, the expertise is encoded by the modulation of specific connectivity patterns. Being manual dexterity a transversal skill that relies on flexible and effective rearrangements of large-scale interactions, α oscillations may represent a suitable candidate for such integrated mechanisms. In fact, this rhythm has been associated with long-range communication, *e.g.* in participants freely engaging in spontaneous behaviors (*e.g.*, reading, watching TV), and coherent α oscillations mediate the communication across large patches of cortex more than other bands ^63^. Then, it is ubiquitous across cortex ^64^.

Conversely, β connectivity seems to be related to the movement itself, thus being highly modulated during both motor tasks, without specific reconfigurations (Figure 2, Figure S2). Based on previous observations, one may expect that the task-induced modulations may reverberate in the successive resting state connectivity, even transiently as in ^65^, through the default rhythm of the sensorimotor system. Infact, in this study, the authors found beta band changes at rest from before to after learning that correlated with motor learning performance, within the hour after. As they suggested, and coherently with our study, these effects may reflect the offline processing of new motor skills. Such an effect is not shown here, as our study is not properly designed to explore the short-term training effects. Rather, we observe a task-induced modulation of the alpha rhythm, the main neurophysiological correlate of the resting brain, likely shaped by long-term manual usage.

### Segregation/integration mechanisms underlying manual dexterity

An apparent dichotomy between stability and flexibility characterizes the architecture of communication in the brain. On the one hand, we expect that such architecture, passing from rest to task, remains stable, as in previous studies ^18, 20–22^. Motor, cognitive, and visual representations are expected to rely on a stable connectivity structure to prevent the degeneration of stored procedural and semantic memories. On the other hand, such a structure must also be flexible to meet the required contextual environment- and task-related demands. Our findings suggest that this balance is encoded in the stability of the spatial topography and the flexibility of connectivity architecture. We observed a stable topography of connections between task and rest (Figure S1). On the contrary, the intensity of the connections and their topology changed depending on the performance (Fig.3, Fig.4A). In the task-relevant connections, see for example between/across MN connections, *high performers* exhibited higher flexibility, *i.e.* stronger changes of coupling strength, while in *low performers* the majority of connections within the motor network were not statistically different between these conditions. The observed flexibility in *high performers* corresponded to specific topological changes related to an increase of functional segregation.

These results are in line with previous studies, where network segregation and integration mechanisms have been associated with behavioral performance ^26, 66, 67^. In our study, in *high performers*, the required segregation seems to steer the system towards an “internal focusing” of task-related networks. In fact, the segregation is observed through an increase of functional modularity, going from rest to task. This agrees with previous studies where modularity has been reported as sensitive to individual differences occurring during motor training ^23, 67, 68^. An increase in modularity during active motor behavior may represent a strategy for responding more efficiently to the task demand. This is supported by Cohen and colleagues ^67^ reporting that increased local, within-network communication is critical for motor execution. Notably, they observed that, during motor execution, the segregation of distinct networks increased (as compared to a working memory task). In line with our results on dexterity, these changes in network organization correlated with better behavioral performance. Our findings suggest that this strategy, based on the increase of segregation and thus modularity, observed in ^67^ between two tasks (motor vs working memory) also applies between task and rest conditions. In fact, learned procedures are automated ^69^ and characterized by focal brain activity ^70^. Increments of network modularity may thus subserve functional specialization, through the selective recruitment of task-dependent networks. To support this idea, the largest rate of reorganization due to connectivity decreases occurred in the motor network, paralleled by a significant decrease of centrality. Specifically, we observed that this increase in modularity is achieved through an important switch of the participation indices of functional hubs (Figure 6). In *high performers,* hubs of the Motor and Control networks showed significant reduction of their participation index when switching from rest to task. The lower integration involving Motor and Control networks, might also suggest that as skill levels increase, individuals no longer make intentional efforts to control the movement, and mechanisms change to become less dependent on volitional movements ^69^ . The observed trend in the participation index suggests a possible shift in the role of the involved hubs from connectors, *i.e.,* linking different communities through local and large-scale connections, to provincial, *i.e.,* short-range connections within the same community or to common nodes. Such a shift seems required to steer the system towards the above-mentioned “internal focusing”. This is in line with previous studies, see for example ^27^. In complete analogy with our findings, Shine and colleagues observed these mechanisms, based on a switch-off of the participation index, when the system moved from an isolated to an integrated state. However, they studied the dynamics of connectivity at rest, *i.e*., in the absence of a task, and thus could not relate this switch to individual performance.

Another important aspect characterizing *high performers* is that, among the strongest connections, representing the minimum set allowing a successful clustering (above the 86^th^ percentile), those involving the Default Mode Network are entirely switched off, when passing from rest to task. This is interesting, since the DMN, typically associated with internal cognition and memory retrieval, is by definition not involved in any task performance ^71^. Furthermore, apart from its own role, the DMN has been consistently reported as a fundamental core of across-network integration ^29, 72, 73^. Thus, switching off any communication with the DMN seems to suggest the decrease of a fundamental axis of integration that seems to be irrelevant for task performance.

As far as it regards *low performers,* we observed an opposite reorganization of the functional architecture, characterized by dysfunctional higher integration. In fact, the loss of network segregation has been reported as a dysfunctional mechanism in many disconnection syndromes, such as stroke, see for example ^74^. Thus, an interesting link seems to exist between functional segregation, efficient behavioral performance, and their decrement during altered task processing. Optimal tuning of the connections implies dense communication among nodes with highly related processing and sparse relationships between areas with different functional roles.

In summary, through the lifespan, learning processes and experience build an intrinsic scaffold of communication that needs to be stable to store procedural and semantic memories. However, this scaffold, embedded in spontaneous activity, in the presence of a task, must also be flexible to enhance and support the individual performance. Our study suggests that in the alpha band, these properties are embedded in the topography and topology of the above scaffold. In particular, the flexibility is realized through specific topological changes that induce an internal focus of task related networks. We found that this pattern of reorganization encodes the individual level of skills. During the task, dexterous individuals reorganize their whole-brain topology, while less dexterous ones have more stable dysfunctional connectivity patterns, especially in task-evoked networks. Notably, this scheme of flexibility/stability is observable in the alpha band, thus a clear marker of manual dexterity.

## Materials and Methods

### Participants

We analyzed Magnetoencephalography (MEG) data from 51 participants (age range 21-35 yo, 25 F, 26 M), a subset of the freely available data collected as part of the Human Connectome Project release (HCP S1200 Release - WU-Minn HCP Consortium). Among the available HCP data, we considered all participants having both rest and task blocks and which satisfied minimal criteria for data quality (see below). HCP data were acquired using protocols approved by the Washington University institutional review board. Informed consent was obtained from all subjects. Anonymized data are publicly available from ConnectomeDB (https://db.humanconnectome.org).

### MEG recordings and experimental paradigm

MEG data were acquired with a MAGNES 3600 scanner system with 248 channels (4D Neuroimaging, San Diego, CA, USA), at a sampling rate of 2034.5 Hz. Subjects were first recorded during three blocks of visual fixation (rest), each lasting 6 minutes, and then during two runs of three types of tasks, the latter of which consisted of a motor task lasting 14 minutes ^32^. The motor task started after about a 10 minute break, during which EMG electrodes were mounted. During the motor task, participants were presented with visual cues providing instructions about the movement to perform with their right or left hand or right and left foot. The hand movements consisted in a finger tapping task involving the thumb and the index finger, whereas for the foot condition they performed toes squeezing. The design of the motor task (Figure 1A) included 32 movement blocks, 8 for each hand and foot lasting 12 seconds and 10 interleaved fixation blocks lasting 15 seconds. Each movement block was composed by a 3 second cue suggesting participants the next movement to perform followed by a 1050 ms black screen period. Then 10 repetitions of visual pacing stimuli lasting 150 ms indicated the beginning of the movement followed by a 1050 ms black screen period in which the movement should be performed. MEG data were recorded together with 4 Electromyography (EMG) channels, placed on each hand and foot, 2 Electrooculography (EOG) channels, and 1 Electrocardiography (ECG) channel. The HCP database also provides pre- processed individual anatomical models computed from structural Magnetic Resonance Imaging (MRI), necessary for source reconstruction.

### MEG data preprocessing and BLP estimation

MEG data released under the WU-Minn HCP project includes unprocessed channel-level signals, channel-level preprocessed and source level processed functional data, together with individual anatomical data. Here, we used channel-level preprocessed resting state data ^32^. Data in the unprocessed format were used for the motor task or when the processed data still contained artifacts. For these data, we applied the same preprocessing pipeline which produced the resting state data, to allow a reliable comparison among conditions. A brief description of the pre- processing pipeline is reported in the following. As a first step, data were band-passed (1.3-150 Hz) and notch-filtered (59-61/119-121 Hz). Then, channels and signal segments contaminated by large artifacts (*i.e.*, excess residual noise in the shielded room or to muscular artifacts) were automatically identified and removed from further analysis. Specifically, noisy channels were identified through the low signal similarity with neighbors, measured through correlation and variance ratio, and through the deviation from the distribution of channel weights obtained by an Independent Component Analysis (ICA)-based approach (using FastICA with deflation approach. Then, we applied again the same ICA approach on sensor space MEG signals to identify environmental, physiological (*e.g.*, cardiac, ocular) and residual channel artifacts, and brain independent components (brain ICs) from sensor space MEG signals, as in ^18, 30, 75^. ICA separation was applied 20 times from different initial conditions. For each of these iterations, the independent components (IC) were automatically classified as brain or artifact. For the classification we used 6 parameters: 1) correlation of the IC Time-Course with those of EOG and ECG channels filtered as the MEG data; 2) correlation of the IC Power-Spectral Density (PSD) with those of EOG and ECG channels; 3) correlation of the IC Power Time-Course (PTC) with those of EOG and ECG channels; 4) temporal kurtosis; 5) 1/f trend of the IC PSD; 6) flatness of the IC PSD. If none of these parameters was above the related thresholds (which are set within the HCP script after a ROC analysis over a set of independent HCP runs), then the IC was flagged as brain IC. The decomposition resulting in the highest number of brain ICs and the lowest level of artifact residual contamination (measured through the average (across the ICs) correlation between the brain ICs and the EOG/ECG channels) was retained as the best iteration and considered for successive steps. In our analysis, the same operator visually inspected the classification of the best iteration for resting state and task data, before proceeding.

The sensor maps of the brain IC were scaled to norm 100 and then projected in the source space by means of a Tikhonov-regularized Minimum-Norm Estimator (MNE). The source space consisted of the individual surface-registered cortical sheet comprising 8004 vertices. The noise level used by the WMNLS estimator was set to 8% of the maximum weight amplitude for each IC. We limited the analysis to a subset of the original 8004 vertices, comprising functionally relevant nodes. Specifically, we considered the parcellation of 164 Region of Interest (ROIs), consisting of vertices belonging to 10 networks depicted in Figure 1B ^73^. We then applied a leakage-correction approach to the IC source-space maps. Leakage is inevitable due to the application of projection schemes to solve the ill-posed inverse problem in MEG. Leakage typically yields a spatially blurred representation of the underlying source distribution. Thus, source-space leakage effects lead to the spurious co[dependence of reconstructed sources, which heavily affects connectivity analysis. Hence, before estimating the functional connectivity, we applied the Geometric Correction Scheme (GCS) ^76^ as in ^19^, where all the related formulas are reported. The GCS is a pairwise approach, *i.e.*, it models and removes the leakage spreading from a source vertex towards all other vertices based on the forward and inverse models. For each seed source, the vector activity of all the other vertices in the set was estimated as the linear combination of the brain IC time courses multiplied by the related leakage-corrected source space weights. For both experimental conditions, we then estimated the Band Limited Power (BLP) time courses as the mean of the activity square module over a sliding window lasting 400 ms, with a sampling rate equal to 50 Hz.

We restricted our analysis to the alpha (8-15 Hz), beta low (15-26 Hz) and beta high (26-35 Hz) bands, filtering the vector activity using separate high-pass and low-pass Butterworth filters. For the motor task only, we also removed the evoked activity before BLP estimation, as in the following. For each direction of the vector activity of each vertex, we first averaged signals over epochs lasting 800 ms. The epoch onsets corresponded to the EMG trigger stored in the HCP database. We then applied the Gram-Schmidt orthogonalization approach as in ^77^, retaining the residual signal for the BLP time course estimation.

### Estimation of BLP Functional Connectivity

The static functional connectivity between a couple of nodes i and j was assessed as follows:

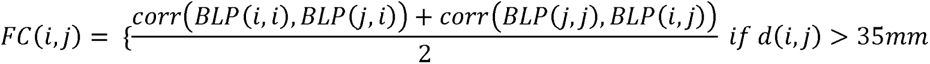

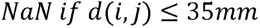

where corr(x,y) is the Pearson’s correlation coefficient between signals x and y, dij is the Euclidean distance between ROIs i and j, and NaN is a “not a number” element for masking correlation coefficients closer than 35 mm. This mask was applied because GCS could be affected by local mis-correction effects, due to seeds mislocalization ^76^. The element BLP (*i,j*) represents the BLP of ROI *i* after removing the bias due to the leakage spread from ROI *j* and the diagonal element BLP(*i,i*) represents the uncorrected BLP of node *i*. We computed the average of the two pairwise correlations to account for slight asymmetries induced by possible numerical errors.

For each run of rest data, the static connectivity was computed over the entire session as the average of the Pearson’s correlation over non-overlapping windows lasting 25 s. For the motor task, the static connectivity was computed over 12 s BLP concatenated segments belonging to the same class of movements. After averaging the interaction matrices across runs, we obtained for each experimental condition individual and group-level static correlation matrices (Fig. 1C). We then computed the z-fisher transform of the correlation data. We then averaged over all the pairwise correlation values involving the nodes in each RSN.

### Analysis of network architecture

To investigate putative changes of the global functional architecture induced by switching from rest to the two motor tasks, the correlation matrices obtained for every subject and experimental condition were analyzed with the Network-Based Statistics (NBS) toolbox separately for each band. NBS is a statistical nonparametric technique that operates directly on raw connectivity values and seeks to identify potential connected structures formed by a set of suprathreshold links (graph components) ^34^. For the comparison between fixation and hands/feet movement, and between *high* and *low performers* (see next subsection), changes in graphs components were tested by using a range of primary (t-statistic) thresholds, ranging from 5 to 7. Permutation testing (n=5000) was then used to ascribe a p-value. Each component identified by NBS satisfied p≤0.05. For the graph visualization, we used the MATLAB toolbox BrainNet Viewer ^78^. For each band, we then counted the relative number of connections changed within the network, across networks and for each network. Then, we analyzed possible links between the correlation changes and behavior. To this aim, according to clustering indices obtained from a k-means algorithm, we split our sample into two groups, comparing the task vs rest differences. Finally, we counted the number of component links modified within and across networks. We are aware that the choice of this threshold certainly influences the size of the obtained components also due to the contribution of false positive links in a component ^34^ . Nevertheless, we here computed the percentage of modified links involving each RSN, normalized by their total number to compensate for the effect of false positive links.

### Regression model task-rest and clustering algorithm

For each subject, band, and network we aimed at estimating at which extent intrinsic connectivity predicted task connectivity. First, for each RSN we considered all within and across-network connections Then, we adopted the following linear model for the task vs rest connectivity for each RSN ^79^:

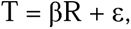

where T is the BLP connectivity for the motor task, R is the BLP connectivity during rest, β is the slope of the linear model, and ε is the error. We used all the pairwise connectivity values of each RSN to estimate β. This provides, for each subject and frequency band, a matrix of β. (network x network). Specifically, β*ij* close to or different from 1 signaled stability/instability between rFC and tFC for the across-RSN interaction involving RSN i and j. Then, we used a data driven approach running a k-means algorithm on the individual beta matrices to cluster them. To estimate the optimal number of clusters, we varied the number of clusters from 2 to 10 choosing the best value as the one producing the maximum average value of the silhouette, provided that its values were always positive (Fig. S3). We then analyzed the link between the clusters and the manual dexterity through t-test comparisons.

To characterize the architecture underlying the identified clusters, we proceeded as follows. For every subject, in the alpha band and for the hand condition, we thresholded the rFC and tFC matrices at the 86^th^ percentile. In this way, we considered the minimum set of strongest connections allowing a successful clustering of *high* vs *low* performers. Then, for each subject, we computed the difference tFC-rFC, we stored separately positive and negative differences, we binarized them and we performed a logical AND across all subjects. In this way, we obtained, for every connection, the percentage of participants in which it consistently decreased or increased. This provided us with a map of the most consistent (we considered connections observed at in 70% of subjects) decreasing or increasing links.

### Analysis of segregation/integration

We then investigated possible links between the individual manual skill and the task-induced modulations of the segregation/integration balance. First, the individual BLP connectivity matrices were transformed into binary graphs according to a percolation analysis ^73^ looking for the maximum threshold ensuring the fully connectedness of the graph (*i.e.*, the number of graph components was equal to the graph size). Then, we applied Louvain modularity, as implemented in BCT (Brain Connectivity Toolbox; ^80^ to estimate the global modularity for each subject and condition (rest, hand motor task). For each subject, the Louvain modularity was estimated 10000 times, retaining the maximum modularity value together with the corresponding modules. We then applied a 2-way ANOVA with condition (Rest, Motor Task) and subject group (according to the clustering results) as factors. Duncan post hoc was applied to significant effects. In addition to segregation analysis, we also investigated how the motor task modulated the central role of networks and hubs. Thus, for each cluster, the nodal participation index (PI) was estimated over the modules in each condition. The PI values were analyzed at the RSN level, through the mean PI over the nodes in each RSN, and at the nodal level. In the first case, we analyzed possible differences in the participation index across groups, conditions and RSN through a 3-way ANOVA. Furthermore, task-induced modulations of the participation index were analyzed through a 2-way ANOVA with RSN (AN, CON, DAN, DMN, FPN, LN, MN, VAN, VFN, VPN) and subject group (according to the clustering results), with Duncan post hoc for the analysis of significant effects. The latter analysis aimed at highlighting opposite trends over groups. Finally, at the nodal level, we ran a t-test comparing task and rest conditions in each group (p<0.05). Nodes significantly changing their participation index were displayed on the cortex through BrainNet Viewer.

## Acknowledgments

This project has received funding from the European Research Council (ERC) under the European Union’s Horizon 2020 research and innovation programme (grant agreement No 759651) to VB.

## Author Contributions

O.M, S.D.P., M.S, A.P., F.d.P., V.B. developed the pre-processing and analysis script. O.M., S.D.P., M.S., A.P., analyzed data. O.M, F.d.P., S.D.P., V.B. drafted the manuscript. S.D.P., & M.C., provided critical revisions. All authors revised the manuscript and approved its final version.

## Supplementary information

### Similarity between rest and motor task BLP correlation

To test the stability of FC topography between rest and task we run a Mantel’s test (Glerean et al., 2016) using 10.000 permutations to calculate a similarity index.

Specifically, in each subject, we measured the correlation between the connectivity submatrices representing each RSN obtained at rest and during the motor task. We found that the two motor tasks preserve the large-scale topography of MEG RSN across bands. This can be appreciated by high correlation values, r>0.86 across conditions and bands (all *p*<0.001), that demonstrate that two motor tasks do not alter the spatial patterns of correlation observed at rest (Fig. S1).

**Fig S1.**
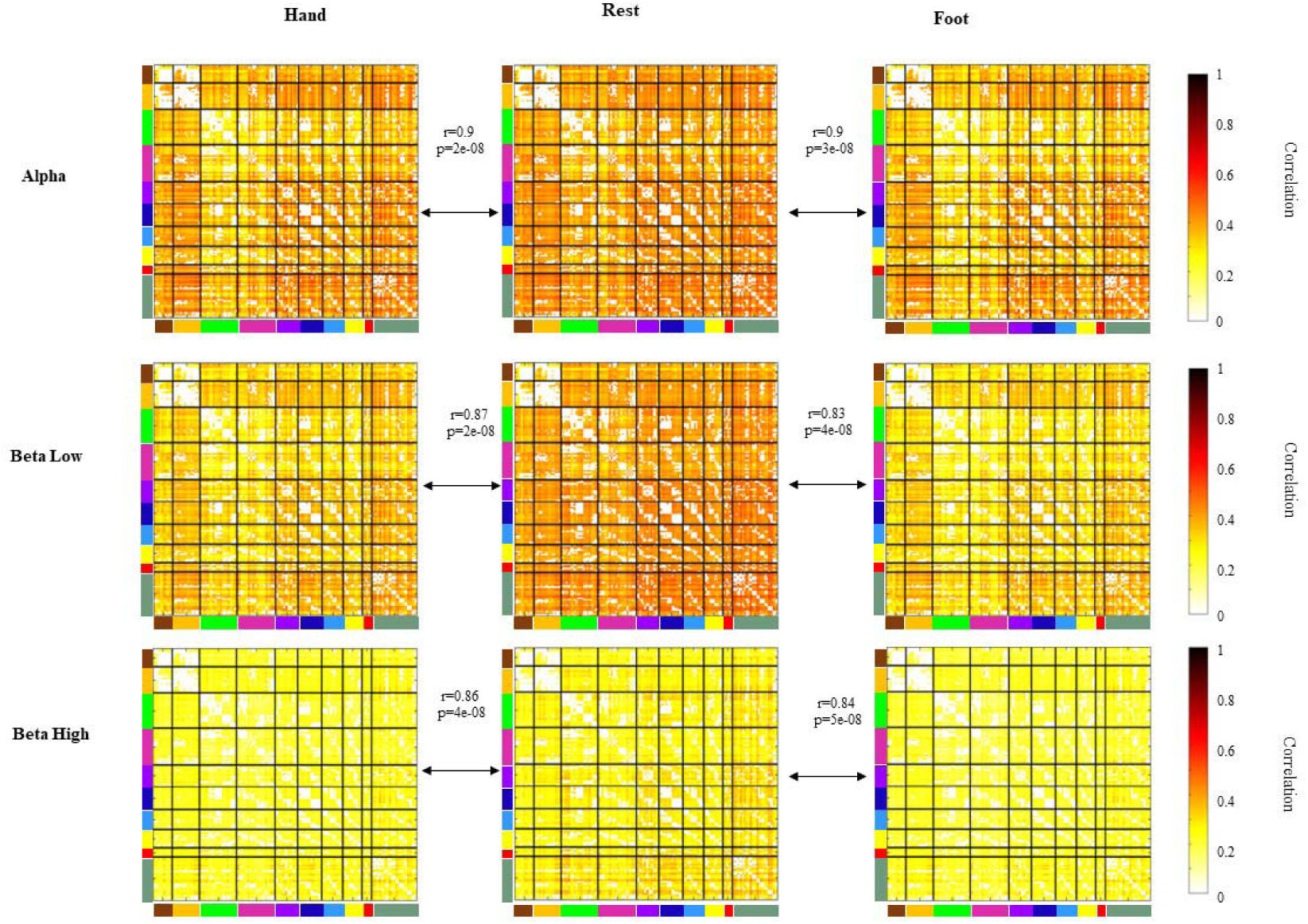
Large-scale topography of connections is preserved across motor tasks and bands. High correlation values (r>0.86) across conditions and bands, demonstrate that the motor tasks do not alter the spatial patterns of correlation observed at rest.

### ANOVA results on modulation of functional connectivity after task performance

To test the modulation of functional connectivity induced by finger tapping compared with toe squeezing, we ran a Repeated Measure ANOVA (RM ANOVA) on the Fisher’s z transformed values of connectivity, with condition (hand, foot), band (α, low β, high β), and network (Auditory network, Control Network, Dorsal Attention Network, Ventral Attention Network, Frontoparietal Network, Language Network, Motor Network, Default Mode Network, Visual Peripheral Network, Visual Foveal Network) as within-subject factors. Results show a main effect of band (*F*_2,100_ = 14.90, *p*<0.0001, pη^2^ = 0.23). Post-hoc comparisons (Bonferroni corrected) revealed smaller decrements of FC in α and larger ones in high beta as compared to all other bands. Specifically, there were significant differences between alpha e low beta band (mean FC_alpha_ =-1.54, FC_betaLow_ = - 3.81; *p*<0.001), between low beta and high beta band (mean FC_betaLow_ = -3.81, mean FC_betaHigh_ =-2.59; *p*<0.05), and between alpha and high beta (mean FC_alpha_ =-1.54, FC_betaHigh_ = -2.59; *p*<0.05).

We found a significant main effect of condition (*F*_1,50_ = 7.01, *p*<0.05, pη^2^ = 0.13) with higher decrements during foot as compared to hand movement (mean FC*_hands_* = -2.31, mean FC*_feet_* = -2.98; *p*_Bonff_ <0.05) (Figg. 2A and S2A). Moreover, we found a main effect of network (*F*_9,450_ = 13.81, *p*<0.0001, pη^2^ = 0.22). The Default Mode Network was the only one showing significant differences with all other networks (except for the Frontoparietal Network) (*p*<0.001) with smaller decrements of connectivity values. The Motor Network (MN) showed higher values of connectivity compared with the Ventral Attention Network (VAN; mean FC_MN_ =-2.61 mean FC_VAN_ = -3.65 *p*<0.01) and lower values compared to the Default Mode Network (DMN; FC_DMN_ =-1.130 *p*<0.001).

**Fig S2.**
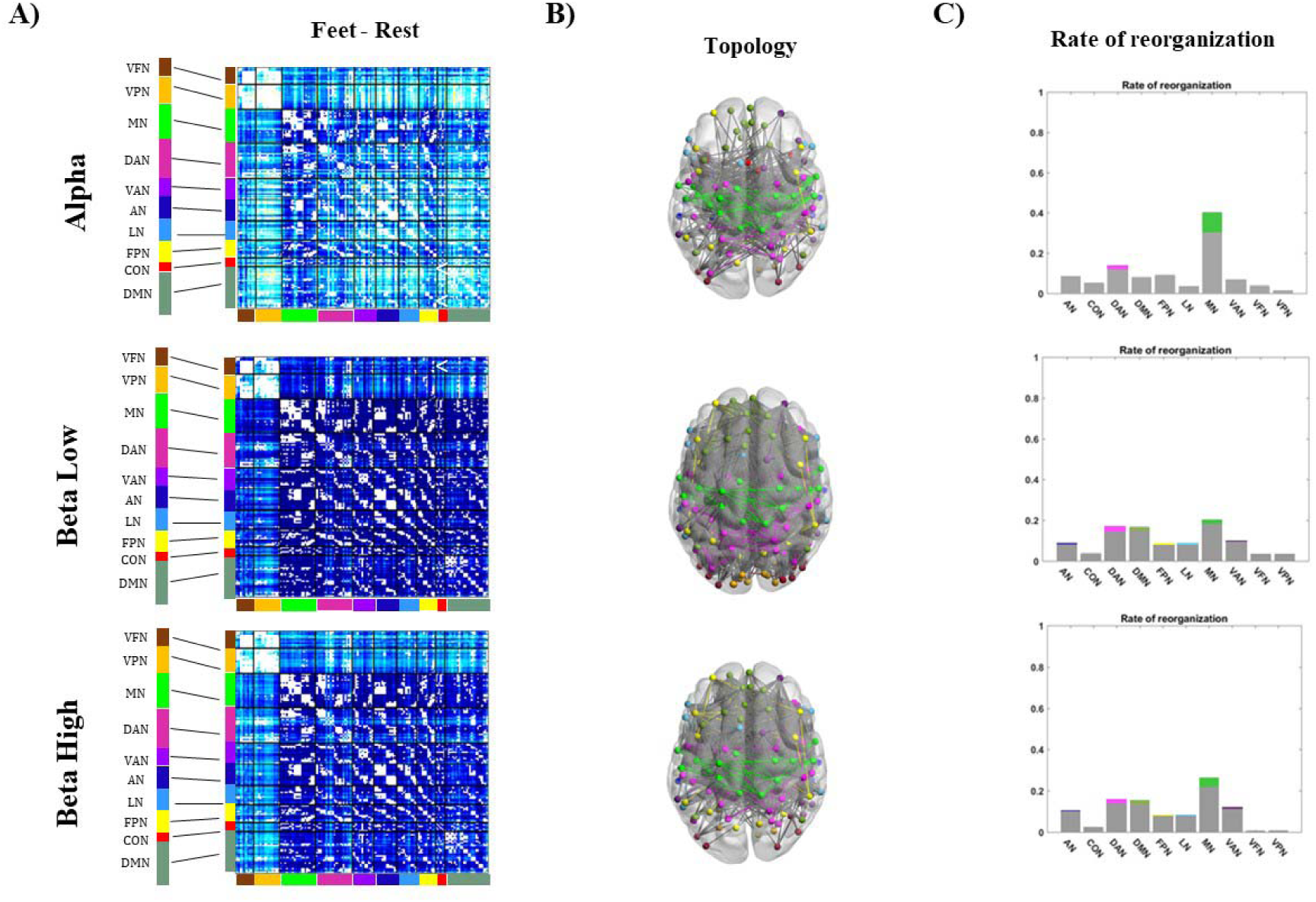
Foot movements decrease the overall connectivity. **A)** Results show a widespread decrease of functional connectivity during toe squeezing in all bands. The decrease during foot movements is higher than during hand movements. **B)** Network Based Statistics analyses show that foot movements lead to an overall brain reorganization larger than with hand movements. **C)** Rate of reorganization in the three frequency bands. Within network connections are color-coded.

Results show a significant interaction band x network (*F*_18,900_ = 9.19, *p*<0.001, pη^2^ = 0.16), with a different modulation of MN in alpha, low and high beta bands (mean FC_alphaMN_ = -1,3, mean FC_betalowMN_ = -3,81, mean FC_betaHighMN_ = -2,72) (all *p_Bonff_*<0.001). FC modulation of all networks in the alpha band was significantly different from network modulations in low beta (higher FC values in alpha compared to beta low; all *p_Bonff_* <0.001). Post hoc comparisons also showed smaller decrements in alpha connectivity compared to beta high for all networks (*p_Bonff_*<0.01) except for the DAN (mean FC_alphaDAN_ = -2.04, mean FC_betaHighDAN_= -2.54, *p_Bonff_* =0.79), VAN (mean FC_alphaVAN_ = - 2.80, mean FC_betaHighVAN_= -3.32, *p_Bonff_* = 0.47) and FPN (mean FC_alphaFPN_ = -1.39, mean FC_betaHighVAN_= -1.47, *p_Bonff_* = 1). We also found a significant interaction effector x network (F9,450 = 3.16, *p*<0.01, pη^2^ = 0.06): in all networks there were higher decrements in FC after foot movements (all *p_Bonff_*<0.001). Finally, band x effector x network effect was significant (F18,900 = 2.68, *p*<0.001, pη^2^ = 0.05).

### Optimization of cluster number for k-means

A k-means algorithm on the beta values matrix was used to test if the data could be clustered in two or more groups. First, we estimated the optimal number of clusters, varying the clusters from 2 to 10 choosing the best value according to the analysis of the silhouette (maximum value of the mean silhouette together with the absence of negative values) (Fig. S3) (Kaufman and Rousseeuw, 1990).

**Figure S3.**
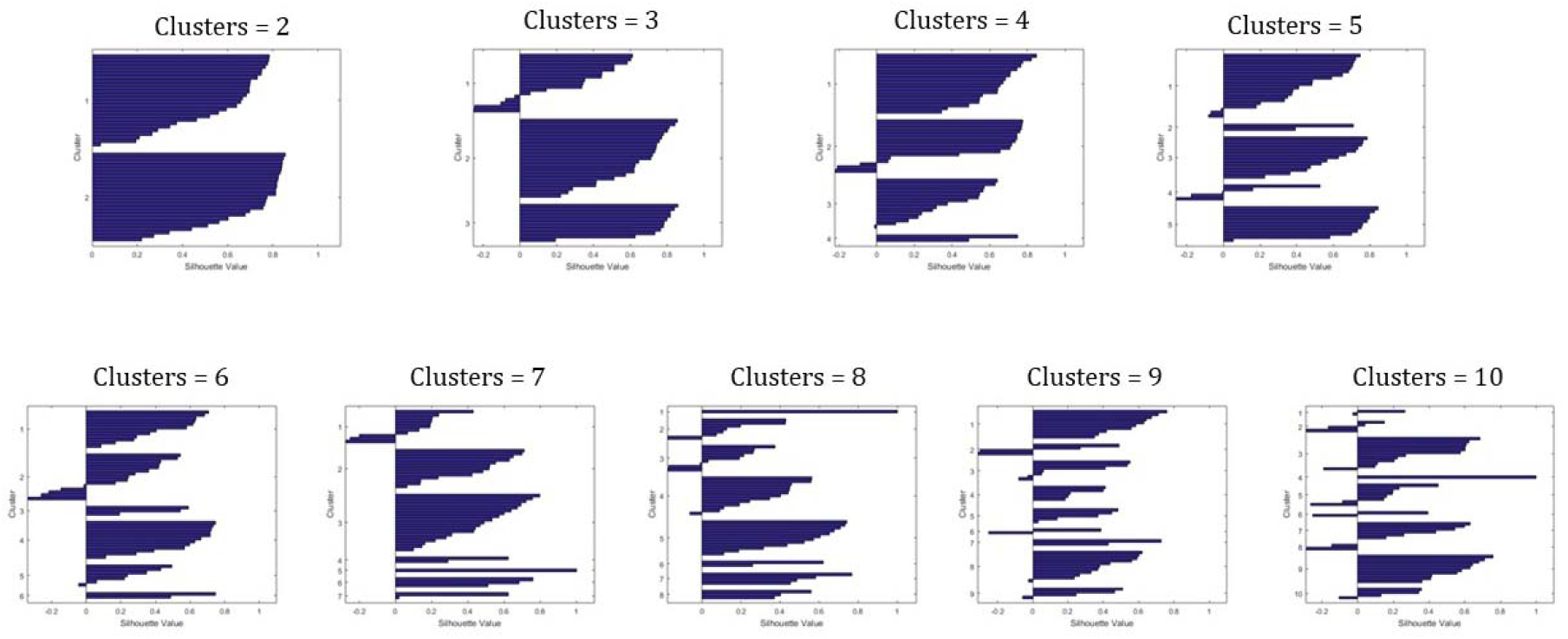
Optimization of the number of K-means classes. Silhouette as a function of the number of classes in the alpha band for right hand. With 2 clusters we obtained the highest average silhouette, with no negative values.

### *Testing the* stability *of the β values and their clustering*

We thresholded the connectivity matrices preserving only the connections above a N percentile value (see Table S1). This corresponds to prune, at increasing level, from N = 0 to N = 0.99, the original set of connections. Then, we computed again the β values for each RSN and we repeated k-means clustering, using the same procedure described above. Our results demonstrate that, within a specific range of percentiles, the algorithm identifies two groups, which differ in dexterity scores. This approach tests the role played by connections of different intensities, *e.g.*, whether the strongest connections mainly drove the classification. Our results demonstrate that the clustering is successful even if we remove the strongest connections (until the 86th percentile).

**Table S1.** **Clustering is effective only in the alpha band.** The numerosity of the classes obtained by K-means at different percentiles (see Materials and Methods) in the alpha and beta bands. No significant effects were obtained in the beta low and beta high, while in alpha the clustering successfully identified high vs low performers at percentiles lower or equal than 0.86.

|  | percentile | group1 (N) | group2 (N) | t-test (p value) |
| --- | --- | --- | --- | --- |
| **Alpha** | 90 | 1 | 50 | .92 |
|  | 85 | 23 | 28 | **.039\*** |
|  | 80 | 26 | 25 | **.015\*** |
|  | 70 | 29 | 22 | **.015\*** |
|  | 65 | 25 | 26 | **.0037\*\*** |
|  | 60 | 23 | 28 | **.016\*** |
|  | 55 | 27 | 24 | **.021\*** |
|  | 50 | 24 | 27 | **.021\*** |
|  | 45 | 27 | 24 | **.021\*** |
|  | 40 | 26 | 25 | **.0059\*\*** |
|  | 30 | 26 | 25 | **.0059\*\*** |
|  | 20 | 27 | 24 | **.0046\*\*** |
|  | 10 | 26 | 25 | **.0049\*\*** |
| **Beta Low** | 90 | 3 | 48 | .085 |
|  | 85 | 48 | 3 | .93 |
|  | 80 | 50 | 1 | .31 |
|  | 70 | 1 | 50 | .31 |
|  | 65 | 1 | 50 | .31 |
|  | 60 | 13 | 38 | .60 |
|  | 55 | 13 | 38 | .60 |
|  | 50 | 37 | 14 | .61 |
|  | 45 | 37 | 14 | .61 |
|  | 40 | 14 | 37 | .61 |
|  | 30 | 16 | 35 | .46 |
|  | 20 | 34 | 17 | .41 |
|  | 10 | 30 | 21 | .33 |
| **Beta High** | 90 | 1 | 50 | .31 |
|  | 85 | 1 | 50 | .31 |
|  | 80 | 1 | 50 | .31 |
|  | 70 | 1 | 50 | .31 |
|  | 65 | 1 | 50 | .31 |
|  | 60 | 1 | 50 | .31 |
|  | 55 | 50 | 1 | .31 |
|  | 50 | 1 | 50 | .31 |
|  | 45 | 1 | 50 | .31 |
|  | 40 | 1 | 50 | .31 |
|  | 30 | 1 | 50 | .31 |
|  | 20 | 1 | 50 | .31 |
|  | 10 | 1 | 50 | .31 |

**Figure S4.**
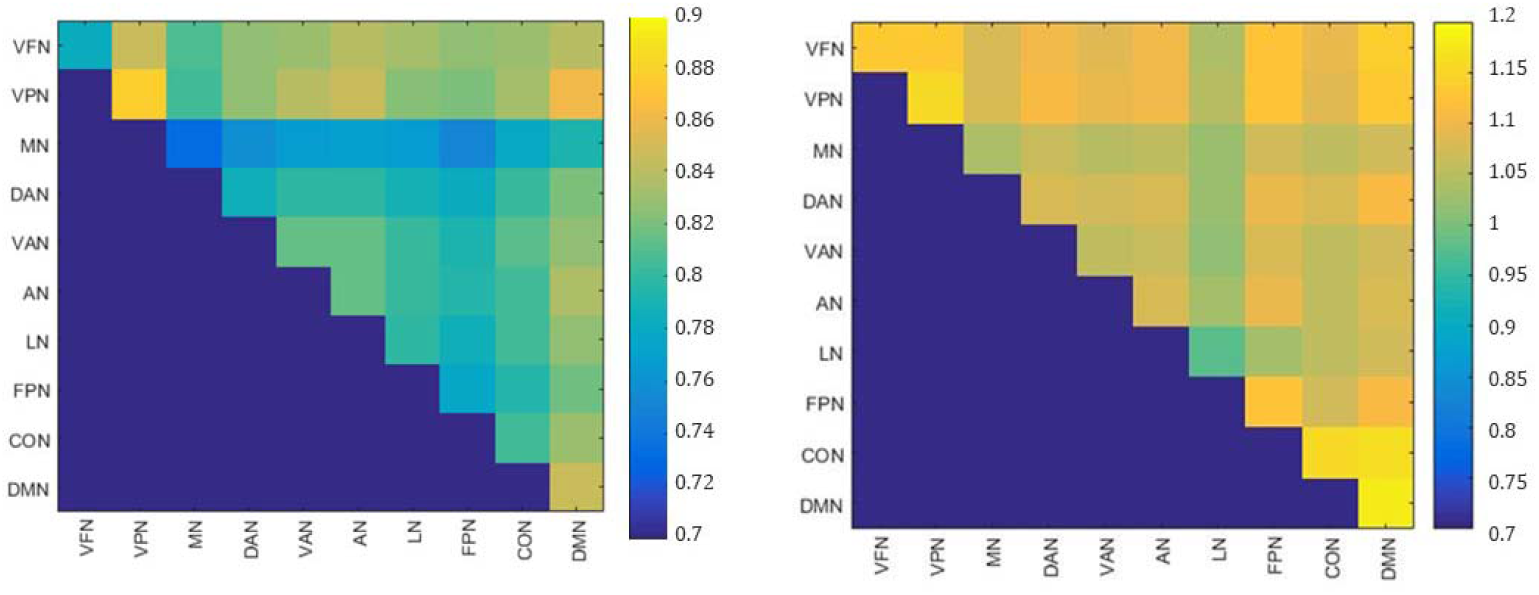
K-means clustering identifies two groups with different beta values profiles. For visualization purposes here we depict beta values matrices with different scales in order to observe how beta changes differently in the two groups.

